# Automated high-throughput heart rate measurement in medaka and zebrafish embryos under physiological conditions

**DOI:** 10.1101/548594

**Authors:** Jakob Gierten, Christian Pylatiuk, Omar Hammouda, Christian Schock, Johannes Stegmaier, Joachim Wittbrodt, Jochen Gehrig, Felix Loosli

**Author notes:** Dr. Felix Loosli Institute of Toxicology and Genetics, Karlsruhe Institute of Technology, Hermann-von-Helmholtz-Platz 1, 76344 Eggenstein-Leopoldshafen, Germany.

## Abstract

**Rationale:** Accurate and efficient quantification of heartbeats in small fish models is an important readout to study cardiovascular biology, disease states and pharmacology at large scale. However, dependence on anesthesia, laborious sample orientation or requirement for fluorescent reporters have hampered the establishment of high-throughput heartbeat analysis.

**Objective:** To overcome these limitations, we aimed to develop a high-throughput assay with automated heart rate scoring in medaka (*Oryzias latipes*) and zebrafish (*Danio rerio*) embryos under physiological conditions designed for genetic screens and drug discovery and validation.

**Methods and Results:** We established an efficient screening assay employing automated label-free heart rate determination of randomly oriented, non-anesthetized specimen in microtiter plates. Automatically acquired bright-field data feeds into an easy-to-use *HeartBeat* software, a MATLAB algorithm with graphical user interface developed for automated quantification of heart rate and rhythm. Sensitivity of the assay and algorithm was demonstrated by profiling heart rates during entire embryonic development. Our analysis pipeline revealed acute temperature changes triggering rapid adaption of heart rates, which has implications for standardization of experimental layout. The approach is scalable and allows scoring of multiple embryos per well resulting in a throughput of >500 embryos per 96-well plate. In a proof of principle screen for compound testing, our assay captured concentration-dependent effects of nifedipine and terfenadine over time.

**Conclusion:** A novel workflow and *HeartBeat* software provide efficient means for reliable and direct quantification of heart rate and rhythm of small fish in a physiological environment. Importantly, confounding factors such as anesthetics or laborious mounting are eliminated. We provide detailed profiles of embryonic heart rate dynamics in medaka and zebrafish as reference for future assay development. Ease of sample handling, automated imaging, physiological conditions and software-assisted analysis now facilitate various large-scale applications ranging from phenotypic screening, interrogation of gene functions to cardiovascular drug development pipelines.

## Introduction

Quantification of heartbeats is an important readout to study physiology and disease states of the heart. The resting heart rate (beats per minute, bpm) is a strong predictive risk factor of overall mortality^1^ and associated with a growing catalogue of genetic variants and environmental-sensitive alleles^2,3^. However, geneticists are facing an enormous challenge to establish causality for associated candidate variants. Small fish models provide efficient means of functional assessment in the context of a vertebrate. In fish models heart rate is examined in biomedical research^4^, including the analysis of inherited or acquired arrhythmia^5,6^, in phenotypic drug discovery and safety pipelines^7^ or in toxicological studies^8^. Moreover, the recent establishment of a vertebrate panel of isogenic strains in medaka fish enables to study variations of quantitative traits such as heart rate in genome-wide association studies^9^. However, there is a lack of efficient screening workflows that allow large-scale quantitative scoring of cardiac phenotypes. Therefore, novel and efficient protocols were needed encompassing sample preparation, automated imaging and image analysis.

The fish species medaka (*Oryzias latipes*) and zebrafish (*Danio rerio*) emerged as major vertebrate models for phenotypic screening, follow-up functional studies, reverse genetics and for studying human pathologies in mechanistical details^10,11^. Medaka and zebrafish have tractable diploid genomes, feature a short generation time (2-3 months), external embryonic development and established protocols for transgenesis and manipulation of gene function including CRISPR/Cas-based approaches^12,13^. Economic husbandry and small transparent embryos are particularly suited for large-scale screening and detailed imaging^14,15^. In both fish species, rapid cardiovascular development leads to a functional cardiovascular system emerging by around 24 hours post fertilization (hpf) in zebrafish^16^ and approximately 48 hpf in medaka^17^. Although the fish heart is two-chambered, its development and function are very similar to the mammalian heart^18^. Strikingly, components of the electrocardiogram of zebrafish and medaka have more similarities to those in humans than have those of rodents, making them ideal models to investigate heart rate and rhythm phenotypes and leverage these properties for drug screens^19-21^. In contrast to mammalian models, early zebrafish embryos are sufficiently supplied by oxygen diffusion allowing the study of severe cardiovascular and dysfunctional hemoglobin phenotypes that result in embryonic lethality in mammals^22^.

Readouts of heart rate and rhythm have been employed increasingly in zebrafish- and medaka-based screens^23-30^. Challenges of current methods to capture heart rate in high-throughput include: (i) immobilization of dechorionated embryos or hatchlings, which in the majority of cases is achieved by anesthesia with tricaine leading to severe adverse cardiovascular effects^31^, (ii) imaging of larvae in standardized orientations, involving labor-intensive manual mounting on dedicated sample carriers or embedding in low-melting agarose, (iii) fluorescent reporter readouts, which preclude studies of specific genetic backgrounds and (iv) lack of user-friendly software for efficient analysis of multiple hearts simultaneously.

We therefore developed a high-throughput screening pipeline with simple sample handling and fully automated imaging of unhatched medaka and zebrafish embryos. Label-free imaging can be performed under physiological conditions and controlled temperature without anesthesia or agarose-based mounting, and with scalable numbers of embryos per plate. The screening workflow is complemented by a user-friendly open-access software package, named *HeartBeat*, for robust quantification of heart rate and rhythm. Applied to both zebrafish and medaka we quantified native heart rate throughout development from heartbeat onset to hatching stage. To facilitate high-throughput applications, microplates containing up to 6 embryos per well were scored with high success rates and reliability. Additionally, we studied cardiac activity responses to temperature ramps enabling the analysis of physical stress response or profiling temperature-sensitive mutants. As a proof of concept for pharmacological screens, we tested nifedipine and terfenadine resulting in heart rate inhibition. This demonstrates the applicability to preclinical drug screens and toxicological tests using fish models.

## Methods

### Fish husbandry and transgenic line

The inbred, isogenic medaka strain Cab and wild type zebrafish AB were kept in closed stocks at the Institute of Toxicology and Genetics (ITG) at the Karlsruhe Institute of Technology (KIT) and the Centre for Organismal Studies (COS) at Heidelberg University as previously described^32,33^. Fish husbandry (husbandry permits AZ35-9185.64/BH KIT, AZ35-9185.64/BH Wittbrodt) was performed in accordance with EU directive 2010/63/EU guidelines as well as the German animal welfare standards (Tierschutzgesetz §11, Abs. 1, Nr. 1). The fish facilities are under the supervision of the local representatives of the animal welfare agency. Embryos were raised at 28°C. Embryonic ages are indicated in hours post fertilization (hpf) or days post fertilization (dpf)^17,34^.

For fluorescent cardiac imaging in medaka, a transgenic line *myl7::eGFP* was generated. A modified plasmid pDestTol2CG (http://tol2kit.genetics.utah.edu/index.php/PDestTol2CG) containing a *myl7::eGFP* cassette was co-injected with Tol2 transposase mRNA into Cab embryos as described^35^.

### Sample preparation and time lapse experiments

One day prior to imaging medaka eggs were optically cleared by rolling the eggs on sandpaper (P1000). One hour before imaging, medaka and zebrafish embryos were transferred from methylene blue-containing medium into plain ERM and E3, respectively. Single medaka/zebrafish embryos were mounted with 150 μl ERM/E3 per well in transparent round (U) bottom 96-well plates (Thermo Fisher Scientific, 268152) using Cell-Saver tips (Biozym). Multi-mounts (≥ 2 embryos/well) were prepared with 300 μl ERM/E3 per well using flat (F) bottom 96-well plates (Greiner, 655101).

For time lapse heart rate measurements microtiter plates were kept between 08:00-20:00 in the incubator at 28°C and transferred to the microscope with 15 min equilibration time before acquiring two loops at 12:00 and 16:00. During night cycle the plate remained in the microscope (constant 28°C) and 4 loops were recorded with 4 h intervals between 20:00-08:00. For all other experiments, medaka embryos were imaged at day 4 post fertilization (start of recordings at 100-104 hpf) and zebrafish embryos at 1.5 days post fertilization (start of recordings at 32-36 hpf) as indicated for each experiment.

### Solutions and drug administration

Terfenadine and nifedipine (Sigma, Steinheim, Germany) were prepared as 50 mmol/L stock solutions in dimethylsulphoxide (DMSO) and stored at –20°C. Prior to application, stock solutions were diluted in ERM (medaka) or E3 (zebrafish) to the desired final concentrations; light sensitive nifedipine was prepared freshly for each experiment. DMSO (vehicle control) and drug incubation experiments were performed with single mounted embryos. For each plate a baseline was recorded in ERM/E3 at 28 °C (see automated microscopy) followed by exchange of the medium with test substances. Prior to the first loop of imaging during drug incubation, embryos were allowed to equilibrate in the incubation chamber of the microscope for min. 15 min to max. 30 min after drug transfer to the plate (equals first point in time).

### Automated microscopy

96-well microtiter plates containing zebrafish or medaka embryos were sealed with gas permeable adhesive foil (4titude, Wotton, UK, 4ti-0516/96). After equilibration for 15 min in the incubation chamber of the microscope under bright-field illumination, the plates were automatically imaged on an ACQUIFER wide-field high content screening microscope equipped with a white LED array for bright-field imaging, a LED fluorescence excitation light source, a sCMOS (2048×2048 pixel) camera, a stationary plate holder, movable optics and a temperature-controlled incubation lid (DITABIS AG, Pforzheim, Germany). The focal plane was detected in the bright-field channel using a software autofocus algorithm. Images were acquired using 130 z-slices (dz=0 μm) with a 2× Plan UW N.A. 0.06 (Nikon, Düsseldorf, Germany). Integration times were fixed with 80% relative LED intensity in bright-field and 100% in GFP-channel with 10 ms exposure time, generating a image sequences of entire microwells of approximately 10 s with 13 fps. Capturing an entire 96-well plate generates 12480 frames resulting in 97.5 GB for full camera sensor readout size. Each microtiter plate was imaged twice at 21°C, 28°C and 35°C or with the indicated time lapse in case of drug exposures.

### *HeartBeat* detection and analysis software

Image optimization was performed prior to the heart rate analysis (Supplemental Material). Based on previous expertise^27^, a new *HeartBeat* software was developed in MATLAB (Mathworks) for embryonic heart rate quantification and segmentation of multiple hearts in a field of view. The *HeartBeat* software features a graphical user interface (GUI) for direct application by non-expert users (detailed instructions for operation see Online Supplement). It is executable with MATLAB runtime (MATLAB 2017) on a Windows computer and available by the authors upon request. We recommend sufficient main memory (≥16 GB RAM) and a solid-state disk hard drive or other storage solution with high I/O rates.

### Data analysis and statistics

Statistical analyses were computed in R language^36^ and data visualized using the ggplot2 package^37^. Differences between samples were tested with Student’s *t*-test and statistical significance accepted at a threshold of *P*<0.05. Multiple comparisons were tested with one-way ANOVA and significant results (*P*<0.05) were analyzed with pairwise comparisons using Student’s *t*-test applying significance levels adjusted with the Bonferroni method. Significant *P*-values are indicated with asterisks (*) with \**P*<0.05, \*\**P*<0.01 and \*\*\**P*<0.001. Correlation analysis was performed using Pearson’s correlation.

## Results

### High-throughput assay and *Heartbeat* software

We implemented a rapid workflow for automated imaging of medaka and zebrafish embryos and a new *Heartbeat* software for extraction of heart rate and rhythm. The multi-step pipeline is schematically summarized in Figure 1. The transparent chorion naturally constrains embryonic movement and therefore supersedes anesthesia or immobilization. This allows simple sample preparation and high-throughput cardiac imaging in a physiological environment. Different speed of embryonic development of both fish species can be leveraged for heartbeat time lapse imaging with different interval lengths ranging between ∼24-72 hpf in zebrafish and ∼2-7 dpf in medaka (Figure 1A). Single or multiple unhatched embryos were analyzed without requirements for positioning, orientation or special mounting medium. We used 96 well-plates as default but also other formats are compatible with the *HeartBeat* software. U-bottom plates are recommended for single embryos per well and F-bottom shapes for multiple embryos per well. A crucial feature of the imaging setup is a stationary plate holder combined with moving optics that eliminate plate movement artefacts and thus obviating mounting media with high viscosity (Figure 1B). Processing with *HeartBeat* is compatible with a wide range of sampling rates, resolutions and illumination modes. Here, default settings for heart rate scoring were 13 fps with 2x detection objective (whole-well detection) and bright-field (Figure 1B). For exact assessment of beat-to-beat variability, higher frame rates are recommended (see *HeartBeat* software).

**Figure 1.**
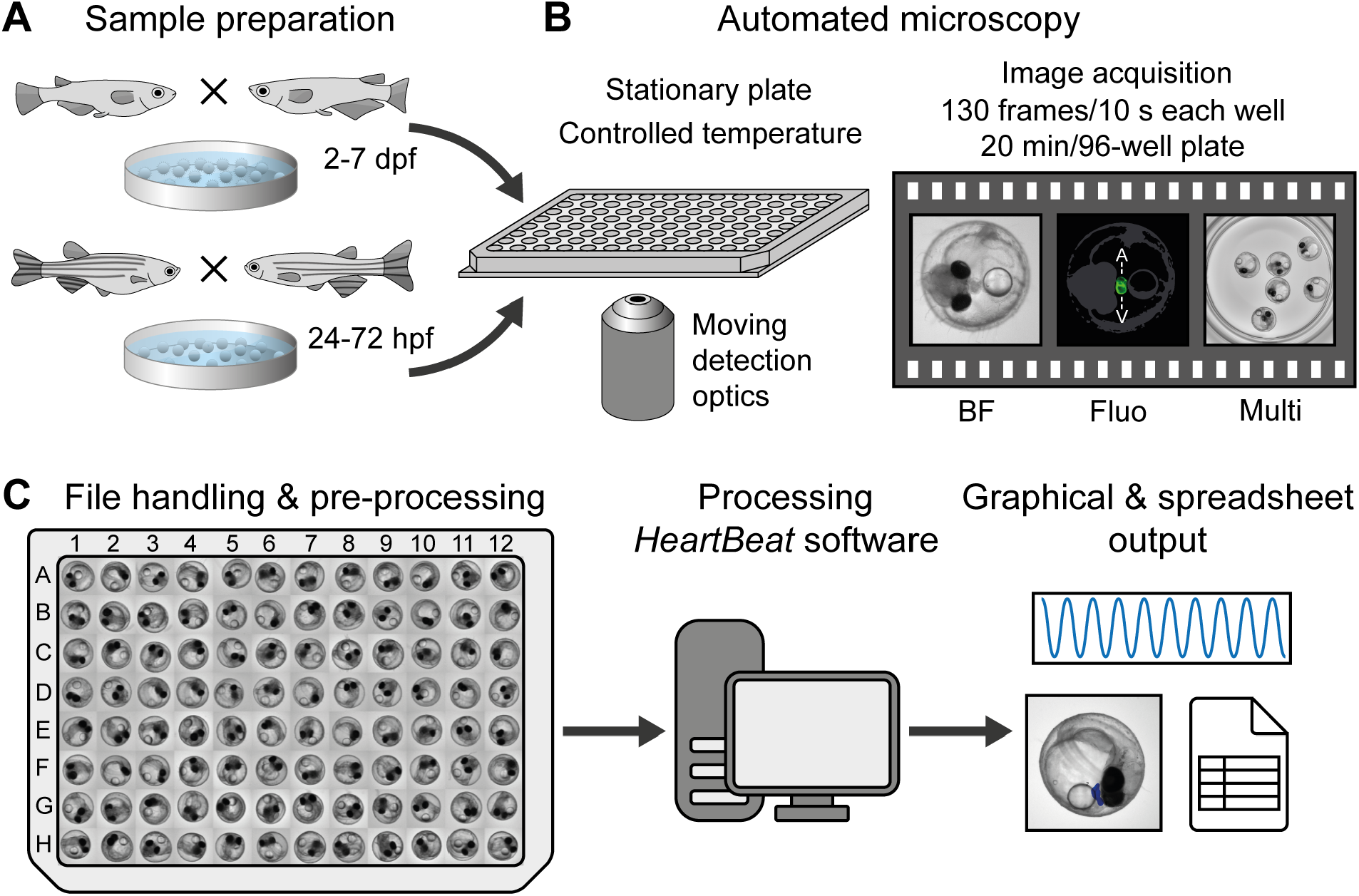
Workflow of automated imaging and heart rate quantification in medaka and zebrafish embryos. **A,** Cardiovascular development is functional after 24 and 48 hpf and hatching occurs around 48-72 hpf and 7-8 dpf in zebrafish and medaka, accordingly, defining time windows of embryonic cardiac imaging in both fish species. **B,** Single or multiple fish embryos per well are mounted without anesthesia or agarose into microtiter plates, which have a fixed position in the imaging platform; image sequences can be sampled at different frame rates, resolutions and channels; all images were recorded with 2x objective, images with single embryos were cropped, multiple embryos in a well (Multi) are presented as full frame; in the fluorescent image A and V denote atrium and ventricle and a rough outline of the embryo is overlaid as a reference. **C,** After image optimization, image sequences are quantified through a GUI of the *HeartBeat* program (compare Figure 2) and heart rates, beat-to-beat variability and an overview of segmentation are saved as spreadsheet and graphical output, respectively.

Whole embryo image sequences are processed by three consecutive operations: (i) segmentation based on cumulative gray value differences in the input image sequence, smoothing and thresholding, (ii) feature extraction by calculating the standard deviation (SD) in identified segments over time, (iii) spectral analysis and classification of heart regions. Dialogue windows ask to specify a range of image sequences in a parent directory and the numbers of embryos contained in each well. Different numbers of fish to be analyzed can be defined for multi-mounted wells. In a GUI, processed time sequences and segments of classified heart regions (blue area) and SD traces are presented (Figure 2A/C) and central settings can be navigated (Figure 2B). Thus, visualization of segmentation and classified heartbeats over time allow for rapid quality check and adjustments. For each image sequence, detailed result files are saved as spreadsheets as well as a graphical overview of the segmentation (Figure 2C). Heartrates of all wells are saved in one summary spreadsheet together with a heatmap (example in Figure 6) in the corresponding parent directory of the dataset. In a sample of *myl7::eGFP* embryos acquired both in eGFP and BF channels, HR was scoreable in 100% in the GFP channel and 98.9% in bright-field (n=92 embryos, data not shown). Heart rate scores inferred from both channels were highly correlated (Pearson correlation coefficient *r*=0.98 for manual count vs. software readout in BF and *r*=0.95 for software readout in GFP vs. software readout in BF; Supplemental Figure 1). In summary, automated heart rate measurements on whole embryo BF images equally perform compared to readout of fluorescently labeled hearts.

**Figure 2.**
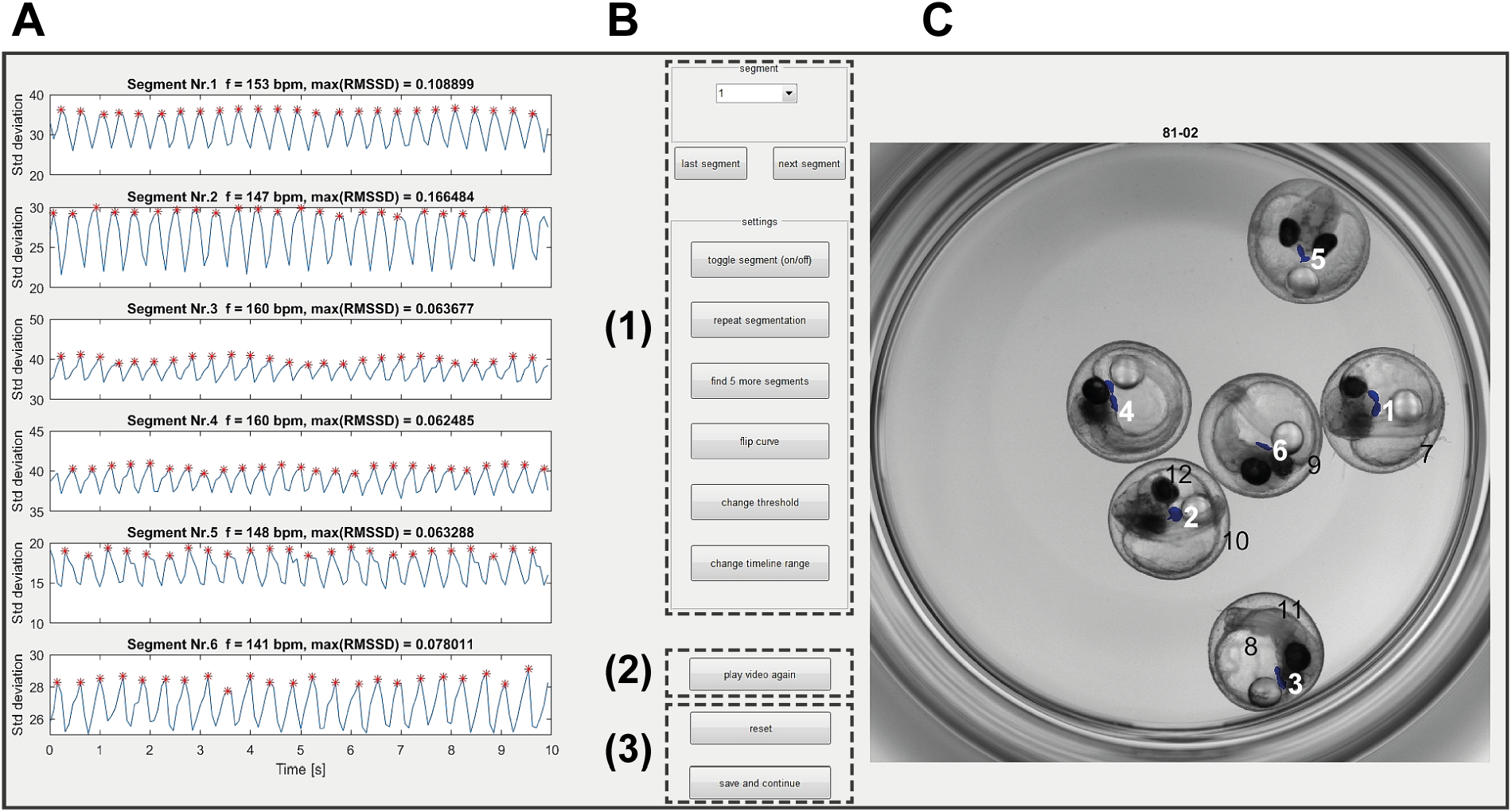
Functionalities of *HeartBeat* software. Graphical user interface of the *Heartbeat* software for computer-assisted semi-automated analysis allowing convenient and rapid quality check. **A,** Standard deviations of pixel values relating to segmented heart regions are displayed over time in the left panel, where heartbeats are indicated by red asterisks for each image sequence. **B,** A control panel allows (1) to refine settings of segmentation and to choose alternative segments, (2) to re-play the current image sequence (visualized in C), (3) to reset all settings or to save heart rates to spreadsheets together with a picture of segmented image sequence. **C,** Visualization of the image sequence with segmented heart regions displayed as numbered blue areas (detailed manual available together with software in online supplements).

### Heartbeat detection of multiple embryos per well

Upscaling the number of embryos per plate can significantly increase throughput for large-scale applications as more specimen can be assayed in a single microplate-wide recording. One option used in some whole-organism screening assays are 384 well plates, that would likewise be compatible with the analytical workflow presented in this study. However, parallel capturing of multiple embryos in a single image, additionally reduces the demand for data storage as well as time of imaging and analysis. For increased throughput, simultaneous readout of multiple fish hearts per imaging field was implemented in the *HeartBeat* algorithm. To assess conditions and efficiency of parallel readout in single wells, an experimental series of 1-6 and 10 embryos per well of 96-well plates were performed for medaka and zebrafish (Figure 3A/B, respectively). Full size image frames were used to analyze multiple embryos with random well positions to achieve correct scoring. Pairwise comparisons of the mean heart rate of mounting groups with ≥2 embryos per well to the reference of 1 embryo per well, yielded for medaka a statistically significant elevation of heart rates in mounts with 2-6 embryos per well and a formally not significant difference for 10 embryos per well well. However, increasing variance and numbers of outliers were observed in wells containing 10 embryos. The equivalent zebrafish experiment (Figure 3B) revealed a higher baseline variability, whereas comparisons to the 1 embryo per well group did not reach statistical significance. A replicate of the experiment for medaka and zebrafish revealed similar results (data not shown). Different offsets of heart rates depending on the number of embryos per well can be controlled by using the same numbers of embryos per well for the entire plate. However, as 10 embryos per well could be scored less effective and yielded an increased number of outliers, a maximum throughput of 6 embryos per well is most efficient and stable. Taken together, the heart rate workflow presented here can be scaled up to 6 embryos per well. For comparability, all wells should be filled with the same number of embryos.

**Figure 3.**
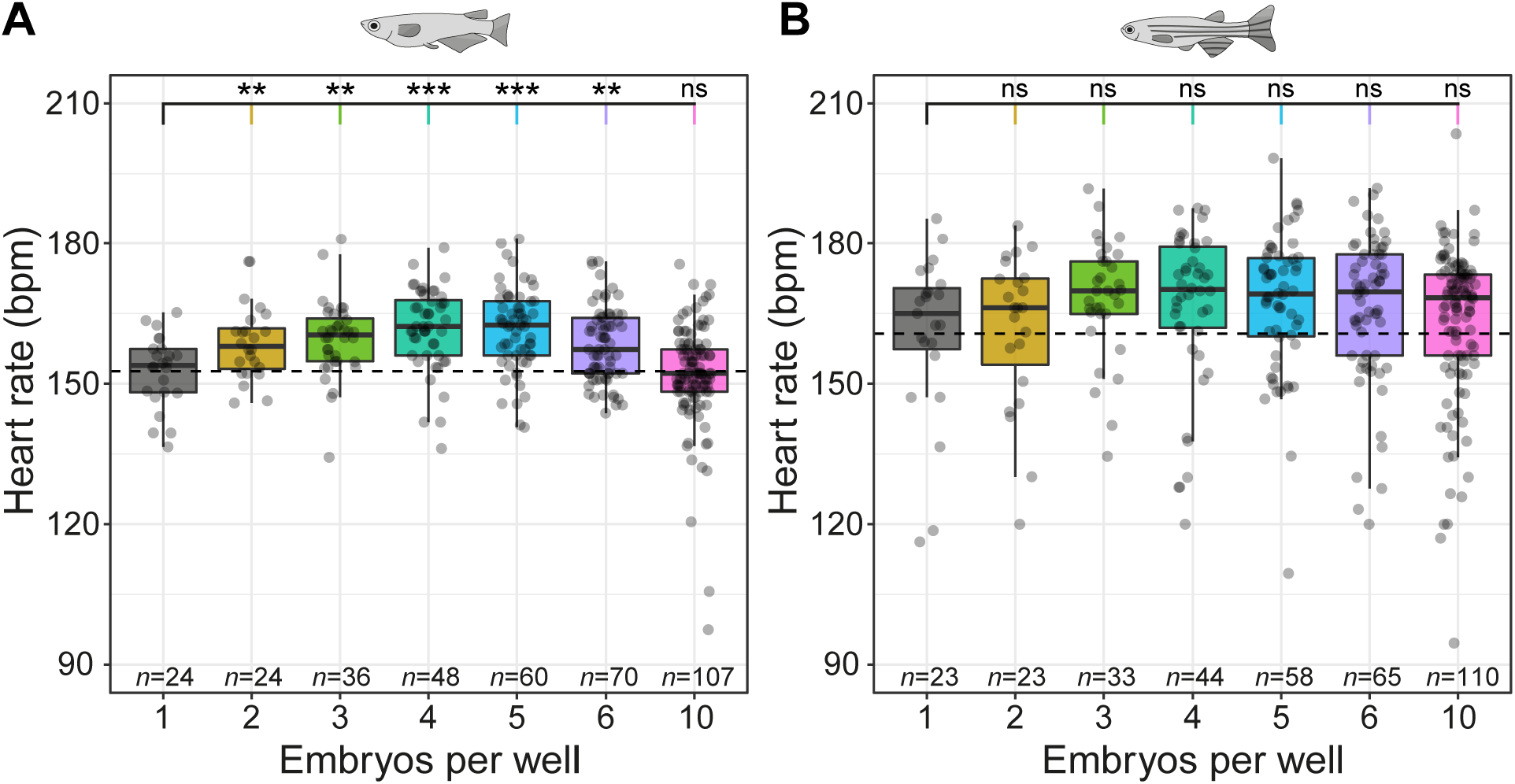
Heartbeat detection of multiple embryos per well. **A, B,** Scoring of heart rate with 1-6 and 10 embryos per well for medaka at 102 hpf (A) and zebrafish at 34 hpf (B). Data is provided as box plots and scatter plots of original heart rate measurements. Significant differences between mounting groups with ≥2 embryos per well each tested against the mean of 1 embryo per well (dashed line) are indicated with \**P*<0.05, *\*\*P*<0.01 and \*\*\**P*<0.001, ns (not significant).

### Heart rate dynamics in medaka and zebrafish development

We applied the assay to score heart rates during the entire development of medaka and zebrafish embryos. Onset of heartbeat varied in the two fish species with earliest heartbeats appearing after 22 hpf in zebrafish and 40 hpf in medaka. From this time points onwards, image data was captured every 4 h (Figure 4). In medaka, embryonic heart rate was rapidly increasing over developmental time and started to oscillate in a day-night rhythm from 72 hpf onwards and reached approximately 175 bpm (at 28°C) around hatching stage (7 dpf, Figure 4A). Similarly, after onset of heartbeat in zebrafish, heart rate increased and plateaued at around 210 bpm (28°C, 72 hpf; Figure 4B). Zebrafish heart rate did not exhibit a day-night dependent fluctuation during the period of analysis (22-62 hpf).

**Figure 4.**
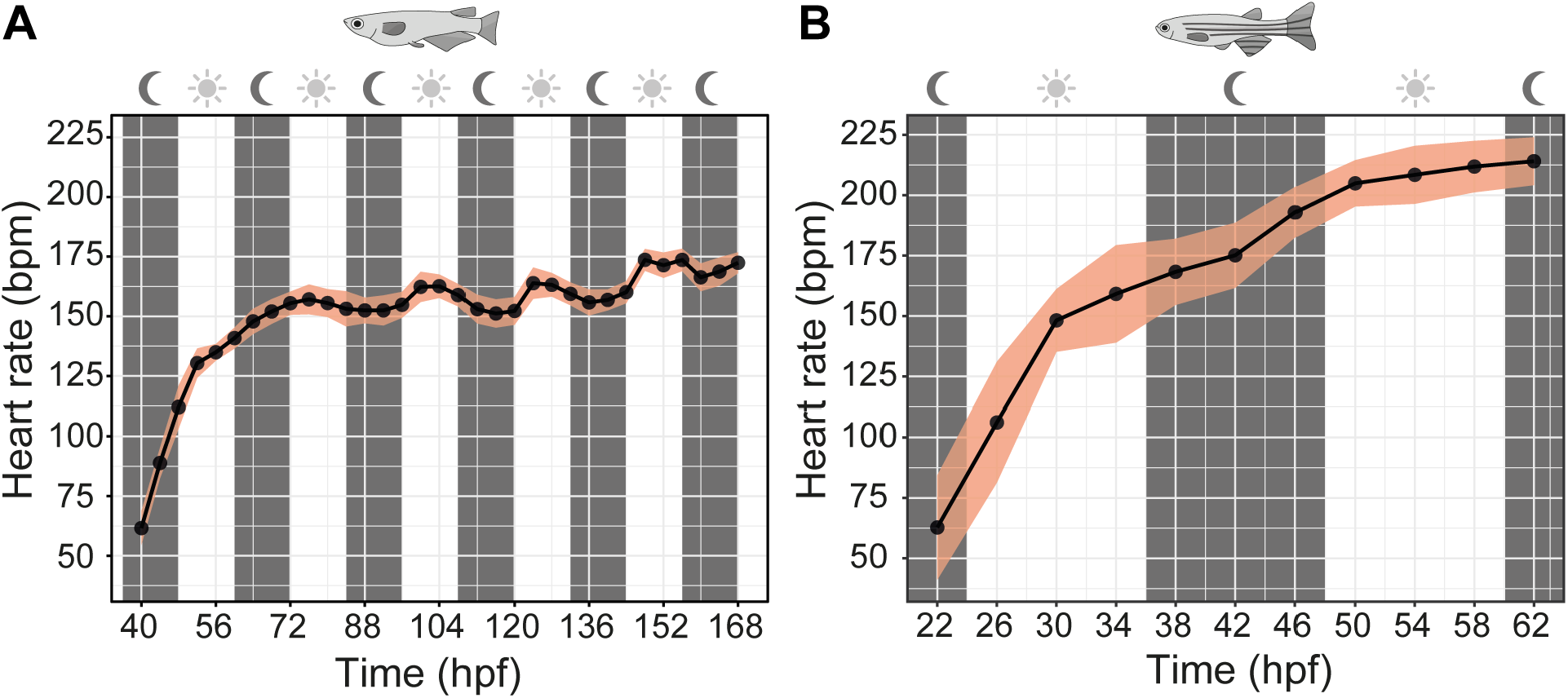
Heart rates of medaka and zebrafish during embryonic development. Embryonic heart rates were measured from onset of heartbeat until hatching stage (details in Methods). **A,** Circadian oscillation of heart rate in medaka from 3 dpf onwards (mean±SD, *n*=13-18 each point in time, total *n*=561). **B,** Heart rate in zebrafish increased continuously and reached a steady state at 48 hpf onwards (mean±SD, *n*=12-52 each point in time, total *n*=396). **C,** Species-specific variability of heart rate during development expressed as SD (derived from data in A and B).

### Temperature-dependent modulation of heart rate

Ambient temperature is a key environmental factor affecting heart rate in most teleosts. To quantify this effect, we studied the influence of ambient temperature changes on heart rate between 21-35°C (Figure 5). Heart rates, both in medaka and zebrafish, accelerate linearly with temperature. Zebrafish exhibited a higher heart rate baseline at all temperatures and a slightly steeper slope of increase (Figure 5B). These results underline the importance of tight temperature control when assessing any heartbeat related cardiac phenotypes.

**Figure 5.**
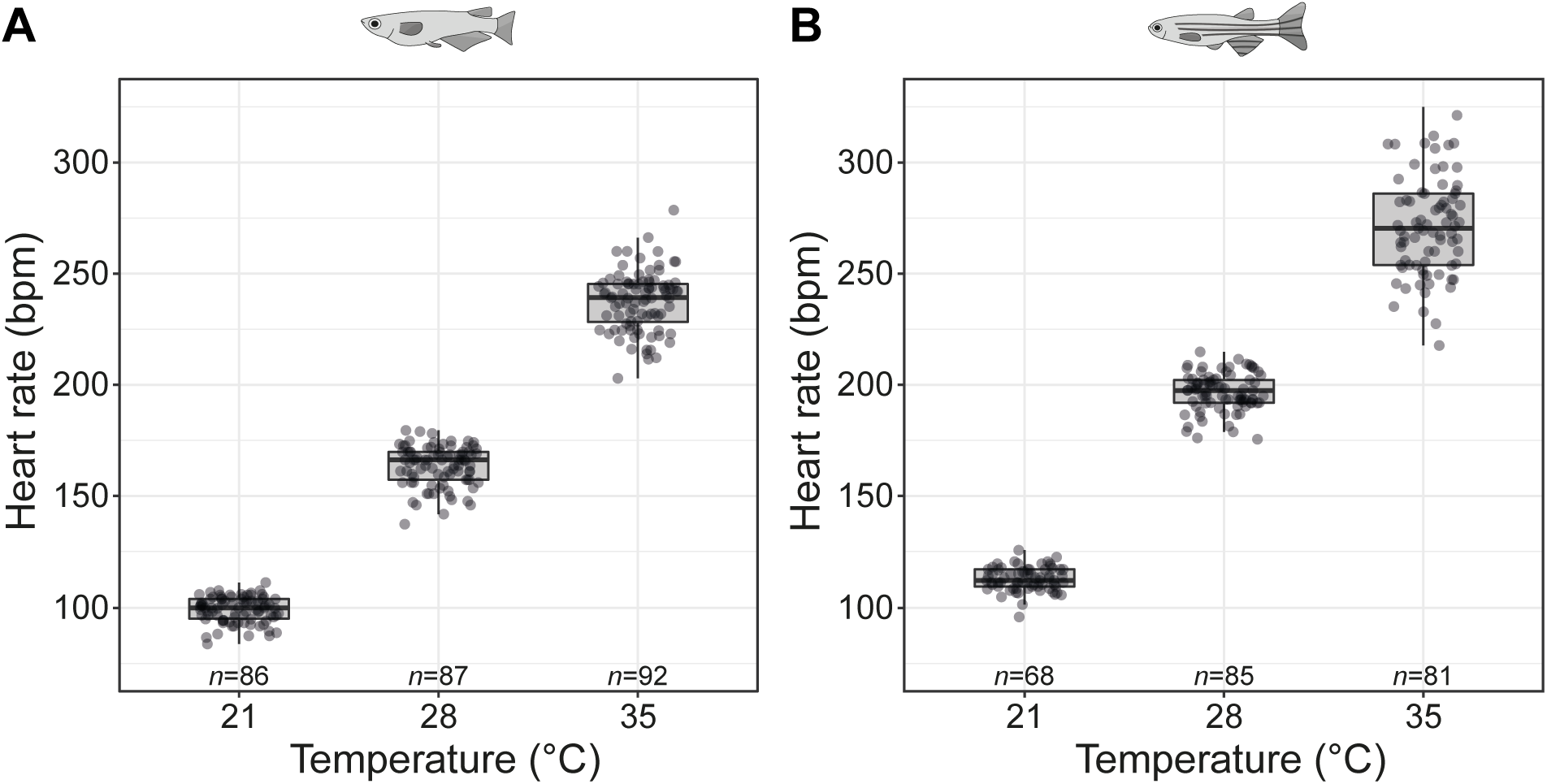
Temperature-dependent heart rate modulation. **A, B,** Heart rate response to temperature ramps in medaka (100-102 hpf, *n*=86-92 at each temperature, A) and zebrafish (36-38 hpf, *n*=68-85 at each temperature, B). Data is given as box plots (median±interquartile range) and overlaid scatter plots of original heart rate values.

### Compound effects on heart rate of unhatched medaka and zebrafish embryos

Heart rate is frequently scored in preclinical screening assays of chemical compounds employing small fish models. To demonstrate the compatibility of our platform with high-throughput chemical screening, we trialed two pharmaceuticals, nifedipine and terfenadine, with known inhibitory effects on heart rate in fish^7,38^. In one 96-well plate, groups of 12 embryos were exposed to terfenadine and nifedipine each at 10, 30 and 100 μM. An internal plate control was kept in embryo medium (EMR/E3). After a baseline measurement, embryos were incubated in the test and vehicle control (0.2% DMSO) solutions for 2 h and captured in 30 min intervals. For each heart rate data set of one microtiter plate, the *HeartBeat* software generates heatmaps for immediate intuitive assessment of drug effects; examples are shown at 90 min drug incubation (Figure 6). Terfenadine elicited significant inhibition of heart rate at 100 μmol/L (μM) both in medaka and zebrafish although with different dynamics. In medaka, 100 μmol/L (μM) terfenadine induced a significant reduction of heart rate after 60 min treatment, which became more pronounced over time (Figure 7A). It induced 2:1 atrioventricular (AV) block in some medaka embryos at 90 min and increasingly after 120 min drug exposure. In zebrafish embryos, 100 μmol/L (μM) terfenadine suppressed heart rate earlier (30 min) and more markedly after 2 h incubation with a transient 2:1 AV block between 30-60 min resulting in dual chamber suppression at 90 min onwards (Figure 7B). Although the current *HeartBeat* software version does not specifically score chamber-specific rates of embryos with AV block (terfenadine), this is nevertheless reflected by atrial or ventricular heartbeats clustering to the upper or lower quartiles of the box plots, respectively (Figure 7A, 100 μmol/L (μM) terfenadine at 90 min and 120 min for medaka; Figure 7B, 100 μmol/L (μM) terfenadine at 30 min and 60 min for zebrafish). Nifedipine inhibited heart rates in medaka embryos equally at 10, 30 and 100 μmol/L (μM) with rapid onset (Figure 7A). In zebrafish embryos, heart rates were decreased by nifedipine in a concentration- and time-dependent manner resulting in asystole for the majority of embryos treated with 100 μmol/L (μM) nifedipine for 120 min (Figure 7B). To provide a reference for embryonic applications using DMSO carrier, heart rates were assessed for a concentration range of 0.2-1%. In medaka, heart rates were not significantly affected by different DMSO concentrations within 2 h of incubation, whereas minor heart rate fluctuations occurred at lower DMSO concentrations in zebrafish (Supplemental Figure 2). As a control for stage-dependent changes of heart rate, all measurements (drugs and DMSO) were normalized to a negative control group of age-matched untreated embryos (Figure 7 and Supplemental Figure 2).

**Figure 6.**
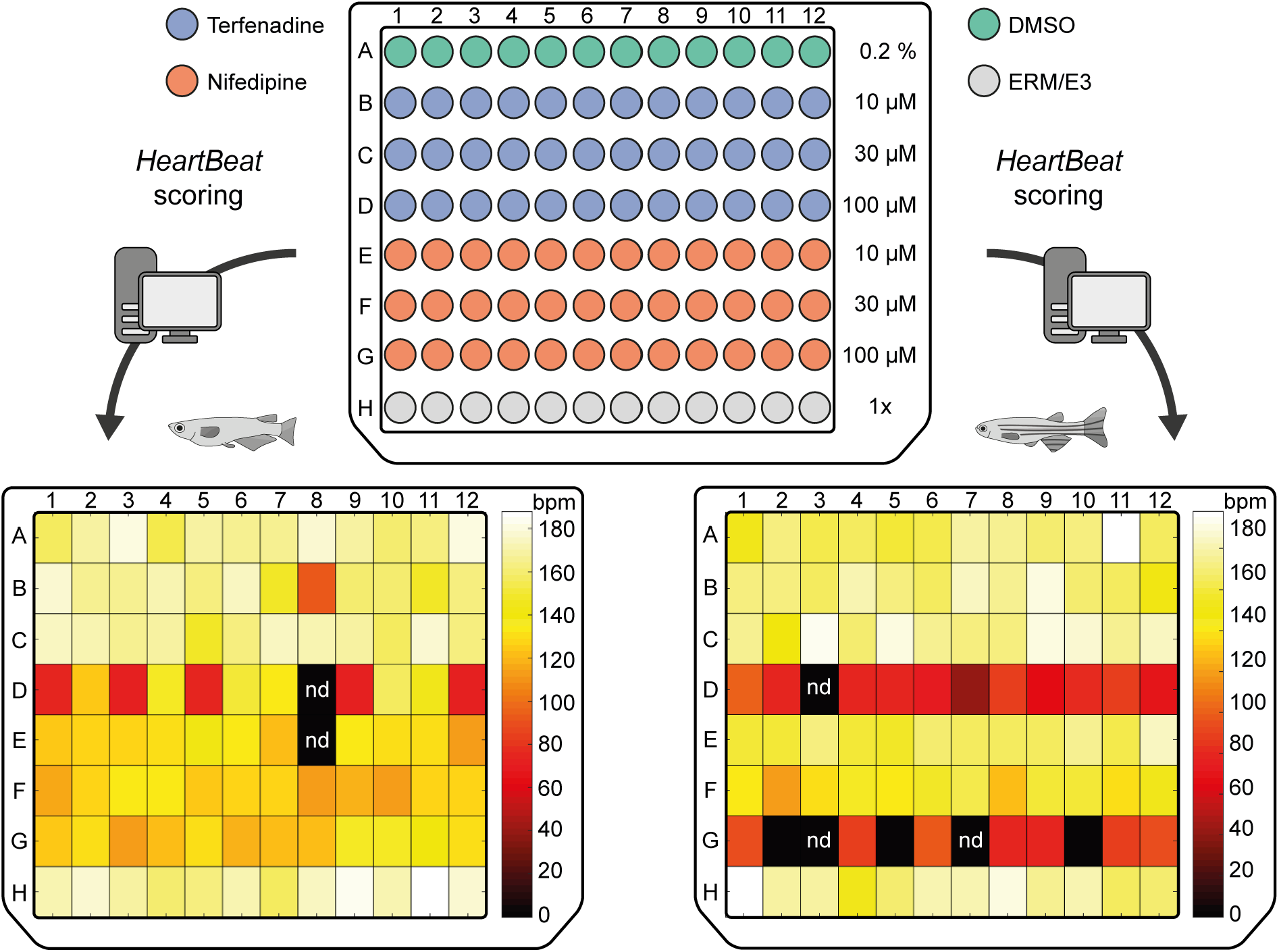
Quantification of heart rates in fish embryos treated with compounds. 96-well format for incubation of medaka and zebrafish embryos with terfenadine and nifedipine each at three different concentrations in μmol/L (μM). As a control for stage-dependent changes of heart rate, all measurements (drugs and DMSO) were normalized to a negative control group of age-matched untreated embryos (row H). The *Heartbeat* software directly provides a heatmap output for intuitive assessment of drug effects (heart rate response shown at 90 min; nd: not detected).

**Figure 7.**
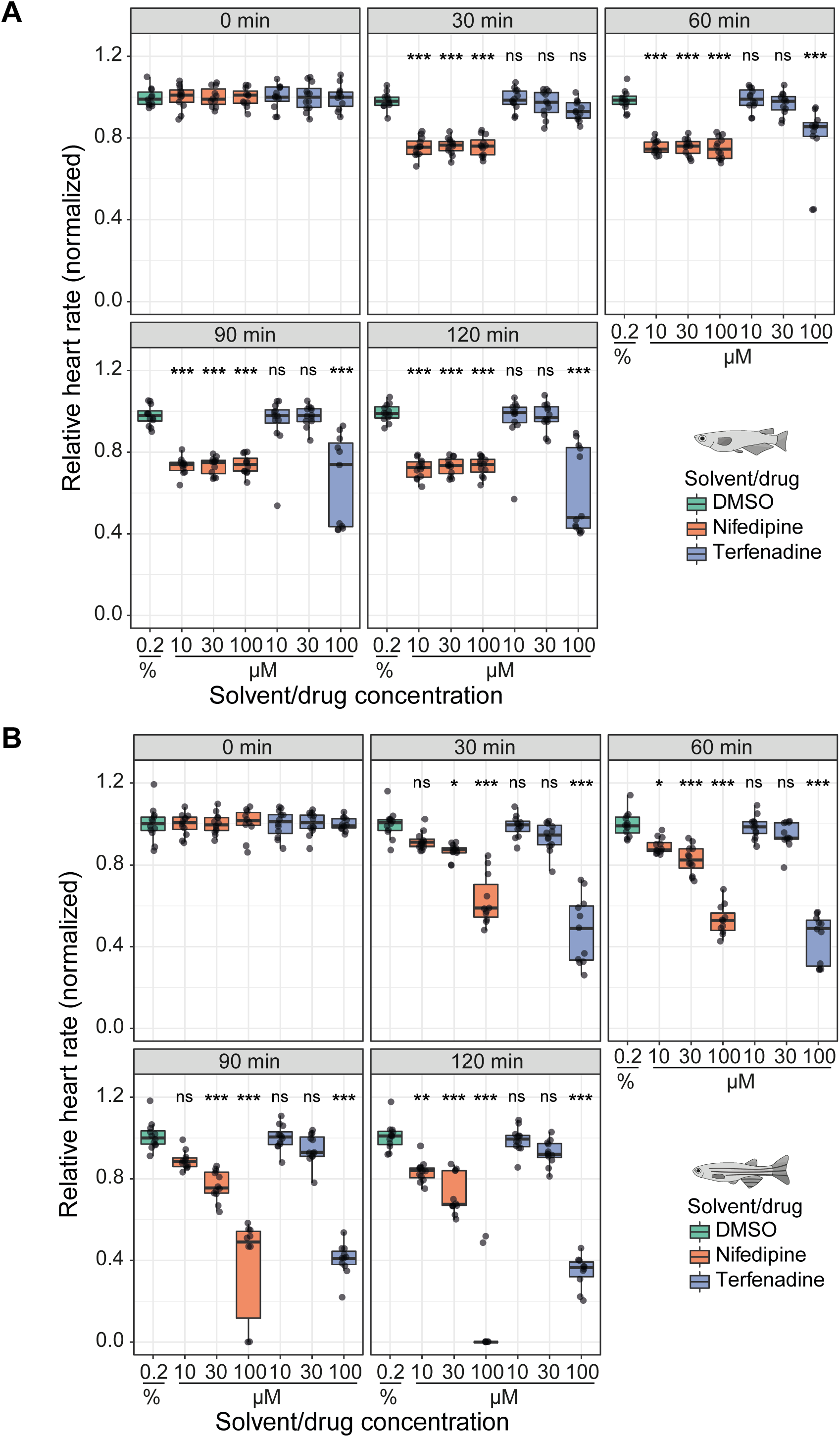
Heart rate inhibition by nifedipine and terfenadine over time. **A,** Time course of drug effects revealing heart rate inhibition in medaka embryos (102-104 hpf) by nifedipine and 100 μmol/L (μM) terfenadine (*n*=11-12 each for DMSO, terfenadine 10/30/100 μmol/L (μM), nifedipine 10/30/100 μmol/L (μM) at every time point). **B,** Concentration-dependent heart rate decrease by nifedipine and terfenadine in zebrafish (32-34 hpf; *n*=10-12 for all treatment and control groups at every time point). Significant difference between each drug concentration and the time-matched DMSO control group were tested with one-way ANOVA and pairwise comparisons using Student’s *t*-test and are given with \**P*<0.05, *\*\*P*<0.01 and \*\*\**P*<0.001, ns (not significant); data is visualized as box plots (median±interquartile range) and scatter plots of original data points. At all points in time, each heart rate measurement is normalized to the corresponding ERM/E3 mean to account for stage-dependent changes and is expressed relative to the corresponding baseline group mean (time point 0).

## Discussion

Heart rate is a key parameter in numerous small fish model applications including genetics and pharmacology and a reliable metric for environmental studies assessing cardiotoxic and adverse developmental effects of small molecules and pollutants. Here, we present a physiological and low-cost assay and open-access *HeartBeat* software for rapid quantitative assessment of heart rate and rhythm of medaka and zebrafish embryos with high analytical power in terms of precision and throughput that permits large-scale phenotypic scoring. The performance of our pipeline allowed to capture concentration-dependent effects of cardiovascular active drugs with high temporal resolution.

Our assay overcomes laborious manual mounting and orientation of samples, motion artifacts, confounding effects of anesthesia and dependency on fluorescent reporters. Fast and reliable quantification of heart rate is achieved by combining automated image acquisition of unhatched, randomly oriented fish embryos with software-based segmentation of heart regions and automatic heartbeat detection. Precise temperature control during automated imaging ensures reproducibility and cross-center data comparability. The *HeartBeat* software was developed for user-independent, automatic segmentation of heart regions and feature extraction of rate and rhythmicity with high reliability and reproducibility. A graphical interface provides non-expert user swift computer-assisted analysis with control of key settings and visualization of analytic output. This interface minimizes user bias and permits to set standards for data comparability between experiments and research groups. A compiled version of the software can be executed with freely available MATLAB Runtime. Of note, heartbeat quantification can be performed robustly on bright-field image sequences with comparable efficiency to image sequences with labeled hearts. The analysis in bright-field mode is key for rapid screening of different strains and precludes fluorescent reporters. The software is compatible with many imaging setups with manual or automated recording using different magnifications. We chose a sampling rate of 13 Hz, sufficiently satisfying Nyquist criterion for frequencies of heartbeats in medaka and zebrafish embryos. Maximal heart rates at 35°C were 278.6 bpm (4.62 Hz) and 325 bpm (5.42 Hz) in medaka and zebrafish, respectively (Figure 5). At 28°C, average heart rates were at 163 bpm (2.72 Hz, medaka) and 197 bpm (3.28 Hz, zebrafish; Figure 5). For analysis of beat-to-beat intervals, however, we recommend sampling at higher frame rates. These frame rates are compatible with standard microscopy equipment. Together with the *HeartBeat* software we provide scripts for file parsing and image optimization, which can be adapted to individual setups and data handling. Cropping to single embryos increased performance, further improved scoring success by eliminating spurious background signals in a storage space efficient manner. TIFF and JPEG compressed image sequences are equally compatible with the *HeartBeat* software. The compatibility with compressed image data massively reduces data volumes, thus facilitating the data transfer across remote collaborators.

To explore the maximal throughput, *HeartBeat* software was applied to readout multiple fish per well and reliably captured heart rates of up to 10 embryos per well. Differences between mean heart rates extracted from wells with different embryo densities (Figure 3) might be reflecting varying physiological environments. For comparative assays this needs to be controlled by using the same numbers of embryos per well for the entire plate. Automated capturing of multiple hearts was performed most efficiently up to 6 embryos per well resulting in >500 embryos processed per 96-well plate, particularly suitable for drug screening pipelines.

To date, several methods have been used to assess heartbeat in zebrafish hatchlings, medaka embryos, fish species, Drosophila, cardiomyocytes or mouse embryos^23,24,26,27,29,30,39-43^. Some of these methods require transgenic embryos with fluorescent cardiomyocytes or erythrocytes to measure heartbeats^28,39^. Other approaches depend on advanced microscopes such as confocal laser scanning microscope^28^ or dual-beam optical reflectometer^41^, imaging modalities that may affect sample temperature and confounding effects on heart rate (Figure 5). Another caveat is immobilization of fish larvae in low-melting agarose to achieve specific orientations, which is time-consuming and complicates genuine high-throughput assays. Some methods were restricted to a narrow time window of newly hatched zebrafish prior to swim bladder inflation to avoid movements of the larvae during sampling^30,39^. In addition, larvae were acclimatized to the illumination of the well scans. To achieve stable imaging condition many protocols rely on anesthesia^23,24,26,29,40^. The most widely used anesthetics is tricaine (MS-222, ethyl 3-aminobenzoate methanesulfonate), that affects cardiovascular functions including heart rate and contractility^44,45^. Our assay, on the other hand, relies on recording non-anesthetized unhatched medaka and zebrafish embryos in their natural environment within the chorion allowing fast sample distribution into microtiter plate, and is compatible with robotic handling of fish embryos in systematic screens^46,47^.

We demonstrate robust heart rate quantification of randomly positioned and oriented fish embryos and implemented options in the *HeartBeat* GUI to easily control artefacts, e.g. through excessive embryo movement during acquisition. Full 96-well plates are imaged in 20 min with 10 s videos at 13 fps for each well. High-confidence quantification can be performed within 45-60 min. In the meantime, our reliable and efficient assay permitted us to automatically quantify heart rate of more than 20 000 image sequences corresponding to approximately 20 TB of data.

While only scant data of embryonic medaka heart rate was available^17^, using the *HeartBeat* assay we now provide a systematic profile as reference for future studies (Figure 4A). Interestingly, after 3 days of development heart rate in medaka showed day-night undulation with a nocturnal dipping resembling the circadian heart rate patterns in healthy humans^48^. Zebrafish has faster embryogenesis and morphological differentiation including looping of the heart is completed by 36 hpf^49^. Serial recordings from onset of heartbeat onwards reliably delineated the rapid increase of heart rate associated with major steps of cardiac development within the first 2.5 dpf (Figure 4B). As aerobic capacity and metabolic rate are directly interrelated with ambient temperature, incremental warming (Figure 5) can be applied as physical stressor in ectothermic medaka and zebrafish embryos, which regulate cardiac output in response to temperature primarily through adaption of heart rate rather than stroke volume^50,51^. Tight control of heart rate by temperature can be leveraged as readout of acute and chronic adaptive stress responses, temperature-sensitive strains^52^ and of susceptibility to arrhythmia in strains with different genetic backgrounds. In summary, our detailed analysis stresses the importance to rigorously control temperature and normalize to stage-dependent changes of beat rates to ensure experimental standardization.

Heartbeat detection is a key metric for drug testing protocols assessing therapeutic and adverse actions of small molecules using fish models. To assess applicability of our assay for compound treatment, we used nifedipine and terfenadine. Concentration-dependent heart rate inhibition and dynamic changes of drug effects were recorded in a 2 h time lapse experiment, reliably correlating with published negative chronotropic effects of terfenadine and nifedipine in zebrafish^7,38^. In human, the antihistamine terfenadine can induce adverse prolongation of QT interval resulting in increased risk for torsades de pointes tachycardia. Detailed studies have demonstrated QT prolongation resulting in AV block in zebrafish^53^ and medaka^52^. Probing effects of terfenadine on zebrafish and medaka embryonic heart rate, distinct time points of induction and time courses of AV block were uncovered by our assay (Figure 7). In zebrafish, nifedipine suppresses cardiac Ca^2+^ channels resulting in a marked decrease of heart rate^38,54^. This drug action was reliably monitored with our assay showing significant effects ranging from sustained inhibition to asystole over time (Figure 7). Taken together, this proof of concept pharmacological experiment in combination with the scalability of our assay raises the prospect of accelerating drug discovery pipelines and rapid pre-clinical prediction of adverse drug effects.

In summary, by combining ease and stability of imaging acquisition with software-assisted analysis our method offers a comprehensive platform for forward and reverse genetics, gene x environment interaction studies, pharmaceutical screens and toxicological assessment, which can efficiently direct downstream studies on mechanistical details of heart rate related phenotypes.

## Acknowledgements

The authors acknowledge the excellent fish husbandry of N. Wolf, N. Kusminski, M. Majewsky, E. Leist and A. Saraceno. Further we like to thank all members of the Wittbrodt laboratory for helpful discussions.

## Non-standard abbreviations and acronyms

dpf: days post fertilization
hpf: hours post fertilization
myl7: regulatory myosin light chain 7
Tol2: Transposon *Oryzias latipes* 2

